# Lowering glucose enhances BACE1 activity and Aβ generation in mouse brain slice cultures

**DOI:** 10.1101/2024.03.12.584616

**Authors:** Olivia Sheppard, Robert Humphrey, Claire S. Durrant, Michael P. Coleman

## Abstract

Numerous environmental risk factors are now recognised as contributors to the onset and progression of Alzheimer’s disease (AD). It is probable that, in most instances, AD arises from a combination of genetic predisposition and environmental influences. In particular, there is a strong correlation between vascular impairment and dementia, yet the specific mechanisms by which vascular impairment and AD are linked, remain unknown. Hypoglycaemia can occur both due to vascular impairment, and due to fluctuating glucose levels in the context of diabetes, another risk factor for AD, and could potentially be involved in disease pathogenesis. To assess whether low glucose could contribute to the build-up of brain amyloid-β (Aβ) seen in AD, we exposed wildtype mouse organotypic hippocampal slice cultures (OHSCs) to varying glucose concentrations. Lowering glucose levels leads to an elevation in both Aβ_1-42_ and Aβ_1-40_ secreted into the culture medium, accompanied by an increased accumulation of Aβ within the slice tissue. This effect is replicated in OHSCs derived from the TgCRND8 mouse model of overexpressed, mutant APP and in human SH-SY5Y cells. The heightened Aβ levels are likely attributed to an upregulation of BACE1 activity, which is also observed with lowered glucose levels. In contrast, OHSCs subject to hypoxia exhibited no alterations in Aβ levels whether singularly, or in combination of hypoglycaemia. Finally, we found that alternative energy sources such as pyruvate, fructose 1,6-bisphosphate, and lactate can alleviate heightened Aβ levels, when given in combination with lowered glucose. This study underscores the capacity to induce an increase in Aβ in a wildtype *ex vivo* system by selectively decreasing glucose levels.

## Introduction

Alzheimer’s disease (AD), the most common form of dementia, is characterised by a decline in memory and executive function. The pathological hallmarks of AD include amyloid-β (Aβ) plaques and tau neurofibrillary tangles (NFTs), which accompany neuron and synapse loss [1–3]. Whilst rare dominantly-inherited genetic mutations directly contribute to increased amyloid precursor protein (APP) expression or Aβ production, the mechanism underlying Aβ and tau aggregation, particularly in sporadic cases which constitute around 95% of AD, remains unclear [4,5]. There are a multitude of partially penetrant genetic and environmental risk factors which are associated with an increased likelihood of developing dementia [6,7]. It is therefore probable that AD is the consequence of a combination of diverse genetic and environmental factors.

Amongst the environmental factors, there is a strong correlation between cardiovascular and cerebrovascular diseases with cognitive impairment and dementia [8–10]. Although a pathological link between impaired vasculature and dementia, including AD, is likely, the specific cause, whether changes in glucose concentration, oxygen levels, or other blood components, remains unidentified. Risk factors for AD often overlap with those for vascular disease, including diabetes, high cholesterol, and an unhealthy diet [11]. Older people with diabetes who have experienced severe hypoglycaemic events are twice as likely to develop dementia [12]. Vascular pathology is prevalent in the vast majority of AD cases [13,14], and a reduction in glucose metabolism is observed in AD brain [15]. Additionally, there is a reduction in glucose transporters GLUT1 and GLUT3 within the hippocampus and cortex in AD [16]. The brain may also be more susceptible to damage when there is a disruption with the vasculature, as it consumes around 20% of the body’s glucose and is the organ with the highest rate of glucose metabolism [17]. Interestingly, Positron Emission Tomography (PET) imaging shows that Aβ deposition tends to occur in areas of the brain with the highest levels of glycolysis [18].

*In vivo* studies indicate that hypoglycaemia can lead to memory impairment, synaptic dysfunction, and an increase in tau phosphorylation in animal models [19,20]. Reduced expression of GLUT1 can also lead to cognitive deficits and neurodegeneration in 6-month-old mice [16,21]. Insulin-induced hypoglycaemia in rats causes increased oxidative stress and microgliosis in the hippocampus [22]. In N2a cells, insulin causes an increase in Aβ trafficking to the extracellular membrane [23]. Hyperinsulinemia in humans is associated with elevated Aβ levels in plasma and cerebrospinal fluid [24]. Conversely, Aβ treatment in rat brain slices makes them more vulnerable to changes in glucose concentration [25]. Moreover, studies have indicated that the toxicity of Aβ can be mitigated by the overexpression of several glycolytic enzymes [26].

Compelling evidence supports the notion that APP and Aβ play a pivotal role in the pathology and progression of Alzheimer’s disease (AD), with subsequent changes such as the aggregation of NFTs and the loss of neurons and synapses. This association is primarily rooted in the fact that dominant mutations linked to familial AD (FAD), found in genes such as APP and the presenilins, tend to exert an influence on Aβ generation, and in particular Aβ_42_ [27,28]. The precise mechanism through which Aβ initiates degeneration remains uncertain, but there are many studies demonstrating its toxicity, particularly in oligomeric form [29–32]. One hypothesis suggests that the aggregation of Aβ, a well as tau, in close proximity to synapses may somehow trigger their loss, ultimately leading to neuronal death [33,34]. However, many plaque-bearing APP transgenic animals fail to show neurodegeneration [35]. Many of these models overexpress human APP with one or several FAD mutations causing an increase in various APP products. However, the vast majority of AD cases are sporadic and lack these mutations, so there is a need for new models which more closely reflect disease pathogenesis in sporadic AD.

The sequential cleavage of APP by β-then γ-secretases gives rise to Aβ, with BACE1 (β-site APP Cleaving Enzyme 1), the β-secretase responsible for APP cleavage, considered the rate-limiting step in its synthesis. Post mortem analysis shows an upregulation of BACE1 in the AD brain [36]. Both oxidative stress and inflammation are characteristics of AD, and are known to increase BACE1 expression and activity [37–39]. Similarly, there are links between oxidative stress and inflammation with hypoglycaemia [40]. BACE1, due to its role in Aβ generation, has become a potential target as a treatment for AD.

Previously, we and others have shown that organotypic hippocampal slice cultures (OHSCs) from postnatal mice are an effective experimental system to probe mechanisms of inflammation, Aβ generation, and tau pathogenic mechanisms [41–46]. To determine whether alterations in glucose concentration could have an AD-related pathological effect upon neuronal tissue *in vitro*, we sought to model hypoglycaemia in wildtype (WT) OHSCs. Here we report that a reduction in glucose in OHSC medium results in an increase in Aβ both within the slice tissue, and that secreted into the culture medium. This occurs in the absence of changes to APP expression, but does correlate with an increase in BACE1 activity. We found no evidence for any alteration in γ-secretase activity, nor any change in transcription of genes for several enzymes associated with amyloid degradation. The low glucose-induced increase in Aβ can be rescued using alternative forms of fuel, including pyruvate, lactate, and fructose 1,6-bisphosphate (FBP). Consequently, our findings lead us to propose that metabolic alterations resulting from glucose availability influence BACE1 activity, and subsequently Aβ production, in this *ex vivo* model.

## Materials and Methods

### Mice

All animal procedures adhered to the regulations and guidelines set forth by the UK Home Office under the Animals (Scientific Procedures) Act 1986, governed by project licenses 70/7620 and P98A03BF9. The research was conducted with the approval of both the Babraham Institute and Cambridge University Animal Welfare and Ethical Review Bodies. Wild-type (WT) animals used in the experiments were sourced from either the C57BL/6Babr colony maintained by the Babraham Institute or the C57BL/6J colony at the University of Cambridge. TgCRND8 mice, characterized by the overexpression of human APP695 with Swedish (K670N/M671L) and Indiana (V717F) mutations [47], were bred as heterozygotes on a 62.5:37.5 SV129:C57BL/6 background, with experiments conducted using littermate controls. For genotyping, DNA was isolated from ear biopsy samples utilizing QuickExtractTM DNA Extraction Solution (Epicentre, QE09050). The tissue was incubated in 20µl QuickExtractTM at 65°C for 15 minutes, followed by vortexing and an additional 2-minute incubation at 95°C. PCR amplification was carried out using the following primers: CCTTTGAATTGAGTCCATCACG and ACAAACGCCAAGCGCCGTGACT . The resulting PCR products were analysed on a 2% agarose gel stained with GelRed®tm nucleic acid gel stain. Both male and female mice were included in the experiments, and all animals were housed in a pathogen-free facility with stringent controls over light and temperature conditions.

### Organotypic hippocampal slice cultures

The generation of organotypic hippocampal slice cultures (OHSCs), was performed as previously described [41,44,48,49]. Briefly, brains were isolated from P6-P9 mouse pups. The brains were bisected along the midline, and sagittally sliced to a thickness of 350 µm using a Leica VT1000S Vibratome. Subsequently, the hippocampus and entorhinal cortex were dissected from the slices and positioned on a fluoro-carbon polymer porous membrane (Millipore, PICM0RG50) with 1 ml of slice culture medium beneath. The composition of the medium was as follows: 50% MEM+ glutaMAXTM (Fisher: 11524496), 23% EBSS (Life Tech: 24010043), 25% Horse serum (Life Tech: 26050-070), 0.65% D-glucose (G5767), 1% Penicillin/Streptomycin (Life Tech: 15140-122), and 6 units/ml Nystatin (Sigma-Aldrich, N1638-100ML). Unless described otherwise, each experiment involved two membranes, each containing three slices from a single pup. This setup allowed for the creation of both a “control membrane” and a “treated membrane” from each individual animal.

### OHSC treatments

Slices were cultured for 13-14 days *in vitro* before undergoing treatment, allowing the initial inflammation resulting from the sectioning process to diminish. OHSCs subjected to glucose reduction were maintained in the same medium, except that glucose-free EBSS was prepared in-house. This custom EBSS solution comprised 1.80mM Calcium Chloride (CaCl_2_) (anhyd.), 0.81mM Magnesium Sulfate (MgSO_4_-7H_2_O), 5.33mM Potassium Chloride (KCl), 26.19mM Sodium Bicarbonate (NaHCO_3_), 117.24mM Sodium Chloride (NaCl), 1.014mM Sodium Phosphate monobasic (NaH_2_PO_4_-H_2_O), and 0.02mM Phenol Red. Hypoxia experiments were carried out using a Binder B170 incubator which for 10% O_2_ treatments, was set to 85% N_2_, 5% CO_2_, and 10% oxygen, and when required, further set to 95% N_2_, 5% CO_2_ and 0% O_2_. Whilst set to 0%, the incubator read at a continuous 0.6% O_2_. The following compounds were also employed: sodium pyruvate (P5280-25G), fructose 1,6-bisphosphate (FBP) (F0752-5G), and sodium lactate (L14500-06). All OHSCs underwent a 7-day treatment period unless otherwise specified.

### SH-SY5Y cells

SH-SY5Y neuroblastoma cells were maintained in DMEM (Themofisher: 41966029) with 10% fetal bovine serum (FBS) (Invitrogen: 10500064) and 1% Penicillin/Streptomycin (Life Tech: 15140-122). Differentiation was carried out using a modified version of Shipley et al. (2016). Cells were plated at a concentration of 50,000 per ml maintenance media. 24 hours after plating, media was changed to DMEM with 2.5% FBS, 1% Penicillin/Streptomycin, 10 µM retinoic acid. Media was replaced after 3 days. After another 3 days, cells were split 1:1 with trypsin and placed in media with DMEM, 1% FBS, 1% P/S, 10 µM RA. 3 days later, cells were placed in differentiation media made as follows: 94% Neurobasal, 1x B-27, 20mM KCl, 1% P/S, 2mM glutamax, 50ng/ml BDNF, and 10 µM retinoic acid. Cells remained in this media until treatments with glucose a week later (with media changes every 2-3 days). For glucose treatment, differentiation media was made with glucose-free neurobasal, and D-glucose (G5767) was added as required. Cells were treated for 5 days and media collected for ELISA.

### ELISA

ELISA analysis was conducted on both culture medium and slice tissue. For medium samples, culture medium was collected before initiating experiments to establish a baseline for each individual membrane. Due to low concentrations of Aβ and APP within the slice, for tissue, OHSCs from multiple pups (number specified in figure legends) were combined, homogenised in 20µl of 5M guanidine HCL, and left shaking at room temperature for 4 hours. Subsequently, 180µl of PBS with protease inhibitors was added before undergoing 20 minutes of centrifugation @ 16,000g @ 4°C. Protein concentration of the supernatant was calculated using BCA assay (Pierce™ BCA Protein Assay Kit, 23225) so equal amounts of lysate could be added to each well. ELISA kits by Invitrogen (mouse Aβ42, KMB3441; mouse Aβ40, KMB3481; human Aβ42, KHB3441; human APP, KHB0051; ultra-sensitive human Aβ42, KHB3544) were used as per manufacturer instructions. OHSC medium samples were diluted 1 in 3 in dilution buffer (supplied with ELISA kit), tissue lysates were diluted 1 in 5, and SH-SY5Y medium diluted 1:2, and each sample was run in duplicate. The plate was read at an absorbance of 450nm against a standard curve (4-parameter fit) using a FLUOstar Omega plate reader.

### Western Blots

Tissue was removed from the membrane using a scalpel and then combined with 100µl of Laemmli buffer (containing 5% SDS, 27.8% glycerol, 138mM Tris at pH 6.8, 0.02% bromophenol blue in H2O), supplemented with protease and phosphatase inhibitors (ThermoFisher: 78442), and stored at -20°C. The samples underwent a further 1:2 dilution in Laemmli buffer (with 10% β-mercaptoethanol) and were boiled for 10 minutes at 100°C prior to loading equal amounts into wells of a 4-20% SDS PAGE gel, which was run at 200V for approximately 1 hour. The protein was then transferred onto a PDVF-FL membrane at 100V for 1 hour. For Dual-color western blotting (used for TgCRND8 lysates as well as their littermate controls), the membrane was immersed in a solution composed of 50% TBS and 50% Odyssey blocking buffer (LI-COR: 927-40003) for 1 hour. Blots were left to incubate overnight on a shaker at 4°C with antibodies appropriately diluted in 5% BSA, 0.05% sodium azide, and TBS-T. Following washes in TBS-T, membranes were exposed to a secondary IRDye anti-mouse or anti-rabbit antibodies (1:10000, LI-COR) for 1 hour in a 1:1 mixture of TBS and Odyssey blocking buffer at room temperature. After an additional washes, the blots were scanned using the LI-COR Odyssey system, and Image Studio Lite 5.2 software was employed for signal quantification. Figures displaying red and/or green bands denote those for which the LI-COR system was utilized. The antibodies used were mouse APP (Sigma: MAB34; 1:1000) and rabbit TUJ1 (Sigma: T2200; 1:2500).

Chemiluminescent westerns were performed on WT tissue due to low levels of APP and BACE1 in an effort to increase sensitivity. Blots were initially blocked for 1 hour in 5% milk/TBS-T. Antibodies were then applied in 5% milk/TBS-T overnight on a shaker at 4°C. Following washes, blots were exposed to goat anti-mouse or goat anti-rabbit secondary antibodies (BIO-RAD) for 2 hours, followed by another set of 3 washes for 10 minutes each. Blots were incubated in either Thermo Scientific™ Pierce ECL Western Blotting Substrate (Thermo: 32106) for 1 minute or Thermo Scientific™ SuperSignal™ West Dura Extended Duration Substrate (Thermo: 34075) for 5 minutes, depending on the strength of the signal. Visualization was carried out using the Amersham™ Imager 680, and ImageJ software was employed for band strength determination. Antibodies for this procedure were rabbit APP (Abcam: ab32136; 1:1000), rabbit TUJ1 (Sigma: T2200; 1:2500), and β-actin (Sigma: A5441; 1:1000).

### Immunostaining

OHSCs attached to culture membranes underwent fixation in 4% paraformaldehyde in PBS for a duration of 20 minutes. Following three PBS washes, the membranes were cut and transferred to a 24-well plate containing PBS. The tissue was then subjected to blocking in a solution consisting of 3% goat serum, 1% Triton, and PBS for one hour at room temperature with continuous shaking. Subsequently, it was incubated overnight at 4°C in the primary antibody diluted in the blocking buffer. Antibodies were used as follows: rabbit TUJ1 (Sigma T2200; 1:500 and rabbit GFAP (Abcam ab7260; 1:1000 ). After three PBS washes, the slices were exposed to the Alexa488 or 568 conjugated secondary antibodies (Life Technologies; 1:250) for 2 hours at room temperature in blocking buffer, followed by another round of PBS washing. Finally, the slices underwent staining with Hoechst (1:5000) in PBS-T for a duration of 10 minutes. The membranes were mounted on slides using Vectashield mounting medium without DAPI (Vector Laboratories: H-1000).

### Quantitative PCR (qPCR)

350µl of RLT lysis buffer was added to tissue scrapped off individual membranes, and RNA was extracted using the RNeasy Extraction Kit in accordance with manufacturer’s instructions (Qiagen: 74104). cDNA synthesis was performed using the Reverse Transcriptase Kit (Quantitect: 205310). The following programme was used on A BIO-RAD CFX96 Real-Time PCR Detection System with c1000 Touch Thermal Cycler: three minutes at 95°C, then 40 cycles of five seconds at 95°C and five seconds at 60°C. Primers are listed in Supplemental Table 1. HPRT, TBP and TUBB3 were used as housekeeping genes. Samples were run in duplicate, and the data is presented using the Delta Delta Cq (ΔΔCq) method.

### BACE1 activity assay

OHSCs were collected in 100 µl of triton buffer (150 mM NaCl, 1% triton x-100, protease inhibitors). BACE1 activity was measured using the β-Secretase (BACE1) Activity Detection Kit (Fluorescent) (Sigma: CS0010). Equal amounts of lysate were added to each well and run in duplicate. A baseline reading was carried out once all reagents were added. The plate was incubated at 37°C for 3.5 hours prior to a final reading. Plates were read using excitation 320 nm and emission 405 nm on a FLUOstar Omega plate reader.

### Statistical Analysis

GraphPad prism software was used for statistical analysis. Statistical tests including student’s t-test and 2way ANOVA were chosen depending on the data set type. Tukey or Sidak multiple comparisons tests were utilized depending on pairing of samples and are specified in figure legends. Results are expressed as mean +/- standard error, and stars represent statistical significance with * = p<0.05, **= p <0.01, *** = p<0.001, ****=p<0.0001.

## Results

### Lowering glucose levels in OHSCs, or a human neuron-like cell line, results in an increase in Aβ

To investigate the potential impact of lowering glucose on Aβ levels in OHSCs, slices obtained from WT mouse pups were subjected to treatment at two weeks *in vitro*. Prior to treatment, medium samples were collected to establish a baseline Aβ concentration. Subsequently, the glucose concentration in the culture medium was decreased from the 40mM used in the standard protocol to 2.8mM. This specific concentration was chosen as it is the product of using glucose-free Earle’s Balanced Salt Solution (EBSS) while excluding the addition of glucose to the regular culture medium.

The glucose levels normally used in slice cultures is considerably higher than in the brain *in vivo* and this can be attributed to the absence of vasculature in the OHSC system and the thickness of the slice tissue (350µm post-cutting). It is presumed that higher concentrations of glucose are necessary outside to facilitate adequate diffusion to slice regions below the surface. Consequently, for the purposes of this study, the term “hypoglycaemia” is tentatively applied to the lower glucose level of 2.8mM. However, it is acknowledged that this concentration is higher than those observed in physiological hypoglycaemic conditions. Glucose concentration within the human brain is reportedly around 1-2.5mM or 1-2µmol/g [50–53], and may drop as low as 0.5mM during hypoglycaemic events [52]. The murine brain is believed to have a typical glucose concentration of approximately 1.2-2.4mM [54,55].

After a 7-day treatment period, medium samples were collected and analysed for Aβ_1-42_ or Aβ_1-40_ concentration using a commercially available ELISA kit. The post-treatment Aβ: baseline Aβ ratio was then calculated to assess the extent of change. For statistical analysis, the treated slices (2.8mM) were compared to the untreated control slices (40mM) from the same animal. Results indicated a significant increase in the amount of Aβ_1-42_ present in the 2.8 mM glucose culture medium compared to control (increase of 40.8 ± 9.5%) (Figure 1A). To ascertain the specificity of this effect to Aβ42, the experiment was repeated using a mouse Aβ_1-40_ ELISA kit. We also found a significant, although smaller, increase in the levels of Aβ_1-40_ in low glucose medium (increase of 20.2 ± 5.7% ) (Figure 1B). Furthermore, we also observed an increase in Aβ_1-42_ in the slice tissue of low glucose treated cultures (increase of 11.7 ± 4.1%) (Figure 1C), although the levels detected in tissue were considerably lower than those seen in medium.

**Figure 1.**
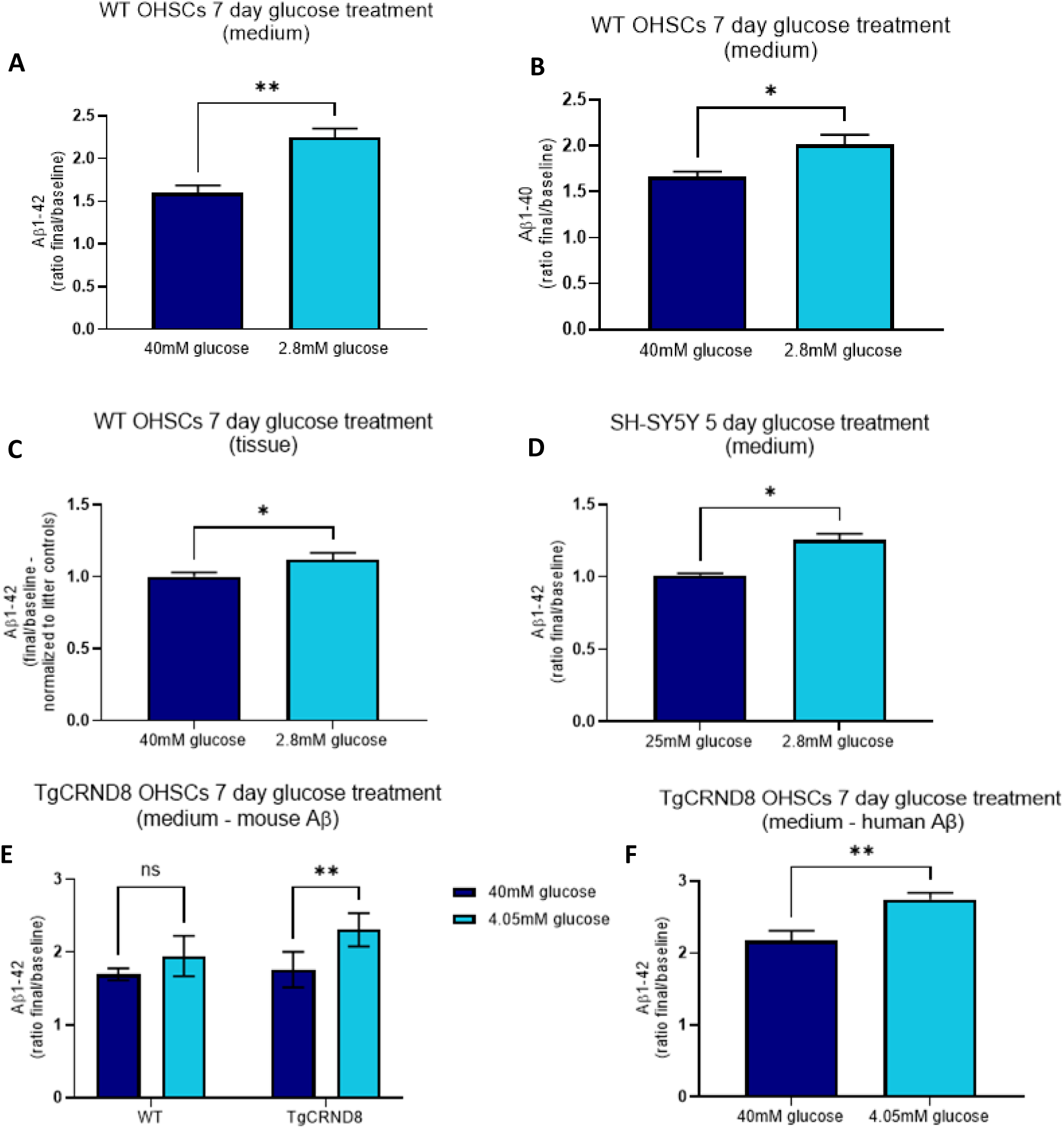
Reducing glucose in OHSC and SH-SY5Y medium results in an increase in Aβ. 14 days in vitro, WT slice cultures had their glucose reduced from 40mM to 2.8mM for 7 days. Tissue was harvested and media collected for ELISA. (a) WT OHSCs showed a significant increase in Aβ_1-42_ (**p=0.0078) and (b) also showed a significant increase in Aβ_40_ (*p=0.014) released into the culture media (n=6; paired t-test). (c) Tissue from WT OHSCs also showed an increase in Aβ_1-42_ levels (*p=0.0199) suggesting the increase is not due to relocation of Aβ from within the tissue into the culture medium, but rather an increase in production, or reduction in degradation (n=10 pooled samples; 31 pups total; paired t-test). (d) To confirm whether a similar effect would occur in human cells, differentiated SH-SY5Y cells were also subjected to a glucose reduction (from 25mM to 2.8mM) for 5 days. There was a significant increase in human Aβ_1-42_ released into the culture media (*p=0.0209) (25mM n=2, 2.8mM n=4; unpaired t-test). (e) TgCRND8 OHSCs exposed to either 40mM or 4.05mM glucose show a similar increase in mouse Aβ_1-42_ released into the culture medium in response to low glucose (**p=0.0045). Their WT littermate controls show a similar trend (p=0.1492). Significant overall effect of glucose treatment (**p=0.0019), no significant effect of genotype (p=0.0864), nor interaction (p=0.1305) (n=5; 2way ANOVA with Sidak’s multiple comparisons test). (f) Human Aβ_1-42_ is also increased in the TgCRND8 culture medium (**p=0.0025) (n=5, paired t-test) suggesting this change is not specific to mouse APP/Aβ. Error bars = mean ± SEM.

In order to determine whether a similar increase could be induced in human cells, differentiated SH-SY5Y cells were subjected to hypoglycaemia. Following medium collection for a baseline reading, SH-SY5Y cells had their glucose concentration reduced from a normal 25mM in their culture medium to 2.5mM for 5 days. The treatment period was shorter than that of OHSCs due to SH-SY5Y cells being unable to survive for a whole 7 days without changing the media. At the end of treatment, the culture medium was collected and an Aβ ELISA performed. Once again, reducing glucose resulted in a 24.6% ± 6.7% increase in Aβ_1-42_ (Figure 1D), suggesting the effect of glucose concentration upon Aβ is not just specific to mouse tissue, but occurs in human cells as well. It is also important to note that this suggests the increase is driven by neurons and does not require the presence of supporting cells such as astrocytes and microglia.

Given that AD likely arises from a combination of genetic and environmental factors, we investigated the impact of a treatment mimicking this environmental risk factor (altered glucose levels) on OHSCs from a transgenic APP mouse line. The objective was to determine whether the effect of this environmental risk factor is additive or potentially synergistic with the genetic risk associated with the transgenic APP. To achieve this, slices from TgCRND8 pups and their WT littermate controls, were cultured in either 40mM glucose (control) or lowered to 4.05mM at 14 days *in vitro*. Despite a more modest reduction in glucose, this was sufficient to induce an increase in both mouse and human Aβ_1-42_ levels in culture medium taken from TgCRND8 OHSCs (Figure 1E-F). There remained a non-significant trend for an increase in mouse Aβ_1-42_ in WT OHSCs. Consequently, the results suggest that the effects of altered glucose and mutant APP overexpression on Aβ are additive.

To confirm that this alteration is not due to an increase in neuronal survival, WT OHSCs were treated with 40mM or 2.8mM for 7 days, or three weeks, and collected for immunocytochemistry or western blot. Immunoblotting with the neuronal marker TUJ1 after 7 days of treatment showed no significant change in protein. Similarly, staining with TUJ1 and GFAP (to determine if there was any difference in astrocyte number) showed no significant differences between treatment after one week (Supplemental Figure 1A-H). However, immunostaining with TUJ1 following three weeks of glucose treatment showed a gross reduction in TUJ1 signal, suggesting neurotoxicity due to hypoglycaemia (Supplemental Figure 1I-J).

### Hypoxia fails to alter Aβ levels in OHSCs

Alterations in glucose and oxygen levels within the brain are typically intertwined due to insufficient perfusion. In an effort to see whether hypoxia could also have an effect upon Aβ both independently, or in combination with hypoglycaemia, WT slices were subject to lowering oxygen. WT OHSCs generated from the same animal were split into four conditions in a repeated measures design: control (20% O_2,_ 40mM glucose), hypoglycaemia (20% O_2_, 2.8mM glucose), hypoxia (10% O_2_, 40mM glucose), and a combination of hypoglycaemia and hypoxia (10% O_2_, 2.8mM glucose). After 7 days treatment, culture medium was collected for ELISA. Although we confirmed that hypoglycaemia leads to an increase in Aβ_1-42_ levels, hypoxia failed to induce any notable effect (Figure 2A). Subsequently, the experiment was repeated with more severe conditions to mimic ischemic episodes. WT OHSCs were subjected to 0mM glucose media in an incubator with 0.6% O_2_ for one hour. Following this, OHSCs were returned to standard culture conditions and left for 7 days, and then culture medium collected for ELISA. Figure 2B illustrates that there was no significant effect on Aβ_1-42_ levels with either treatment.

**Figure 2.**
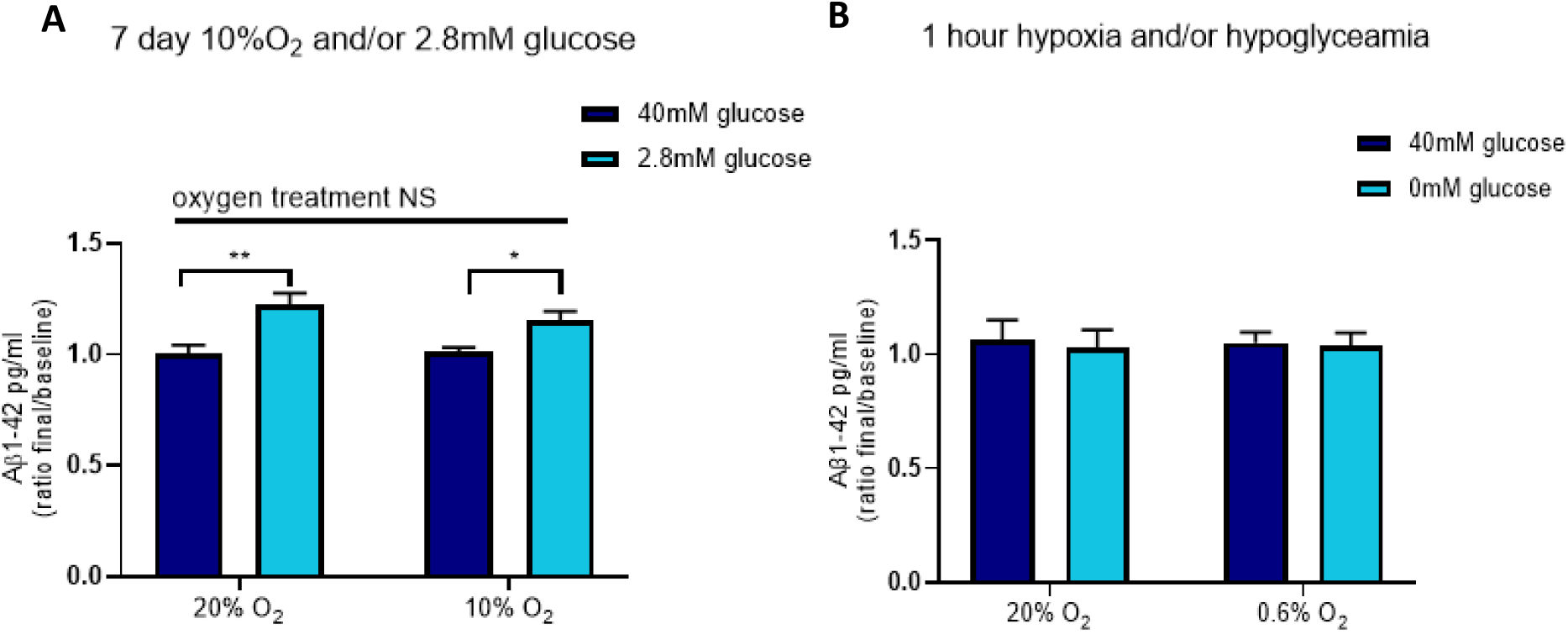
Hypoxia does not alter Aβ_1-42_ levels in OHSCs. (a) WT OHSCs were subjected to a reduction in glucose from 40mM to 2.8mM, this time in combination with a reduction in oxygen from 20% O_2_ (normal atmospheric oxygen) to 10% O_2_ for 7 days. There was once again an effect upon Aβ_1-42_ due to a reduction in glucose regardless of whether cultures were in 20% or 10% O_2_ (p=0.0013 and p=0.0018 respectively), but no hypoxia dependent change in 40mM or 2.8mM glucose (p=0.9936 and p=0.2768 respectively). There is a significant overall effect of glucose (***p=0.0005), but not O_2_ (p=0.6350), nor glucose: O_2_ interaction (p=0.1551) (n=9; 2way ANOVA with Tukey’s multiple comparisons test). (b) Slice cultures were subject to more severe conditions mirroring an ischemia event. Glucose was reduced to 0mM and oxygen was decreased to 0.6% O_2_ for 1 hour. OHSCs were then placed in normal culture conditions for 7 days prior to collecting media. This resulted in no change in Aβ_1-42_ levels (p=0.9846 for O_2_ and p=0.6090 for glucose) (n=5, 2way ANOVA). Error bars = mean ± SEM.

### Glucose-induced increase in Aβ is not associated with altered APP transcription or translation

The low glucose-induced increase in Aβ could potentially be due to an increase in APP transcription. However, this scenario seems unlikely. As depicted in Figure 1D, the elevation of human Aβ is observed in glucose treated TgCRND8 slices, where the human APP transgene is regulated by the prion promoter [47]. This implies that whether the promoter is prion-based or the typical mouse promoter for APP, a comparable increase in Aβ secretion is observed in response to alterations in glucose levels. To confirm, mRNA was extracted from WT OHSCs after 7 days of glucose treatment, and qPCR was conducted. The results revealed no significant increase in APP mRNA (Figure 3A).

**Figure 3.**
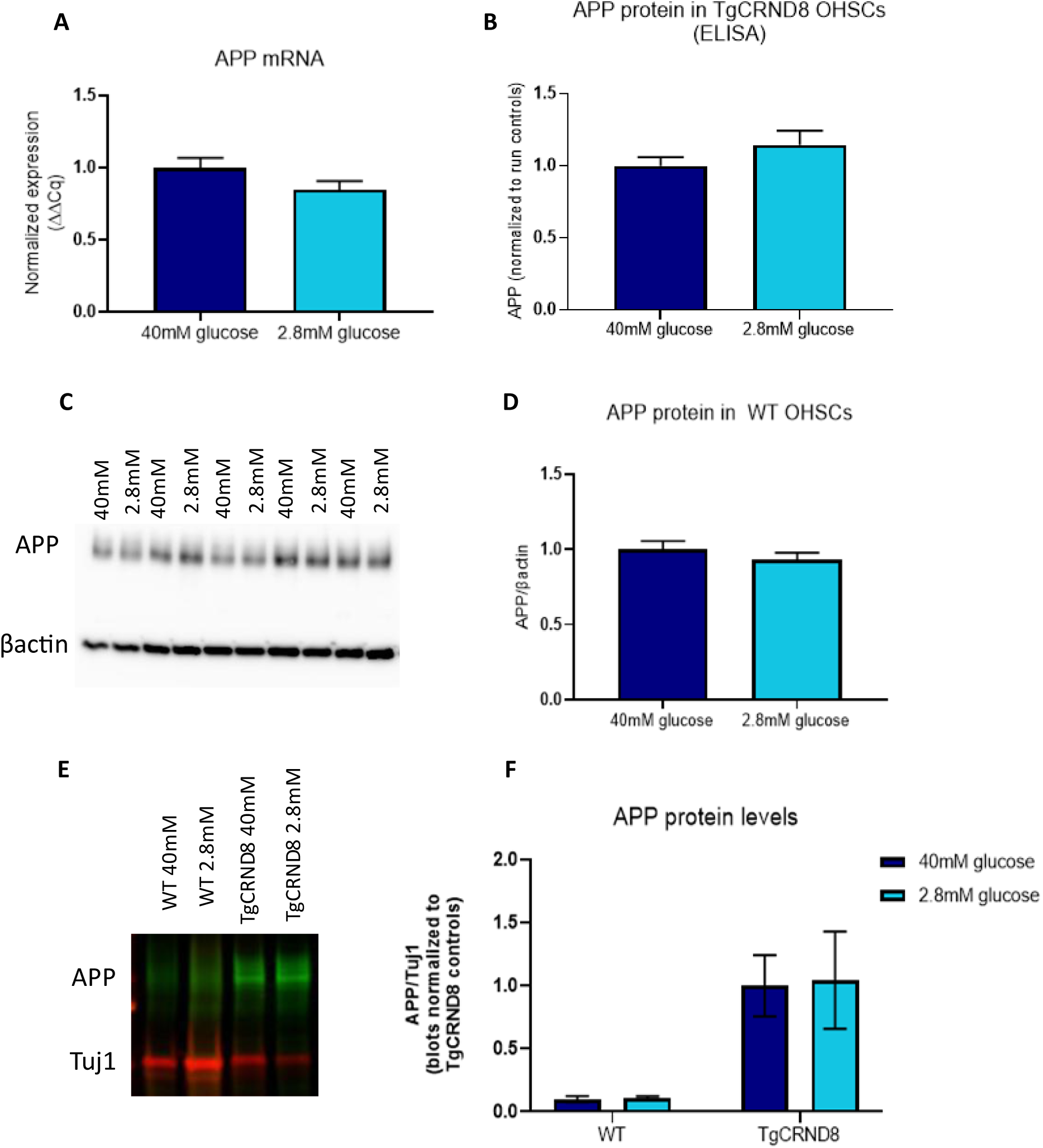
Hypoglycaemia-induced change in Aβ is not due to a change in APP mRNA or protein. (a) WT OHCSs subjected to a reduction in glucose from 40mM to 2.8mM for 7 days and had their tissue harvested for RNA isolation. qPCR showed no evidence for an increase in APP mRNA (p=0.0903) (n=12, paired t-test). (b) Following treatment, TgCRND8 slices were pooled and collected for APP ELISA which showed no significant effect upon APP (p=0.1427) (n=6 pooled samples; 16 pups total; paired t-test). (c-d) Western blot analysis of APP protein in WT tissue also shows no significant change (p=0.3009) (n=13, paired t-test). (e-f) Similarly, western blot showed no change in APP protein in TgCRND8 lysates, nor their littermate controls (WT p=0.9998; TgCRND8 p=0.9930). No significant overall significant effect was seen due to glucose treatment (p=0.9473), nor genotype (p=0.0512) (WT n=2; TgCRND8 n=5; 2way ANOVA with Sidak’s multiple comparisons test). Error bars = mean ± SEM.

Once determining that there was no evidence for a change in APP transcription, changes in APP translation were investigated. After subjecting slices to glucose treatment for 7 days, TgCRND8 OHSC tissue was collected for ELISA. Pooled samples, each containing tissue from three or four pups, were collected per sample to ensure an adequate amount of tissue for the assay. No significant increase in human APP protein levels was identified by ELISA (Figure 3B). Similarly, tissue from both WT (Figure 3C-D) and TgCRND8 (and their WT littermates) (Figure 3E-F) OHSCs were used for immunoblotting, and showed no significant change in APP protein.

### Hypoglycaemia results in an increase in β-secretase activity

In the amyloidogenic pathway, APP undergoes sequential cleavage by the β- and then γ-secretase enzymes, ultimately leading to the production of Aβ. To investigate the mechanism behind the observed increase in Aβ production following hypoglycaemia, the activity of the β-secretase BACE1, was analysed using a commercially available BACE1 activity kit. WT OHSCs were subjected to control or low glucose treatment for 7 days and then collected in triton buffer. We found a significant increase in BACE1 activity in OHSCs exposed to low glucose conditions (Figure 4A). Subsequent examination using qPCR to assess transcription revealed no significant effect on BACE1 mRNA (Figure 4B). Western blotting demonstrated a slight decrease in the amount of BACE1 protein in the low glucose slices (Figure 4C-D), potentially a compensatory response to the increase in activity.

**Figure 4.**
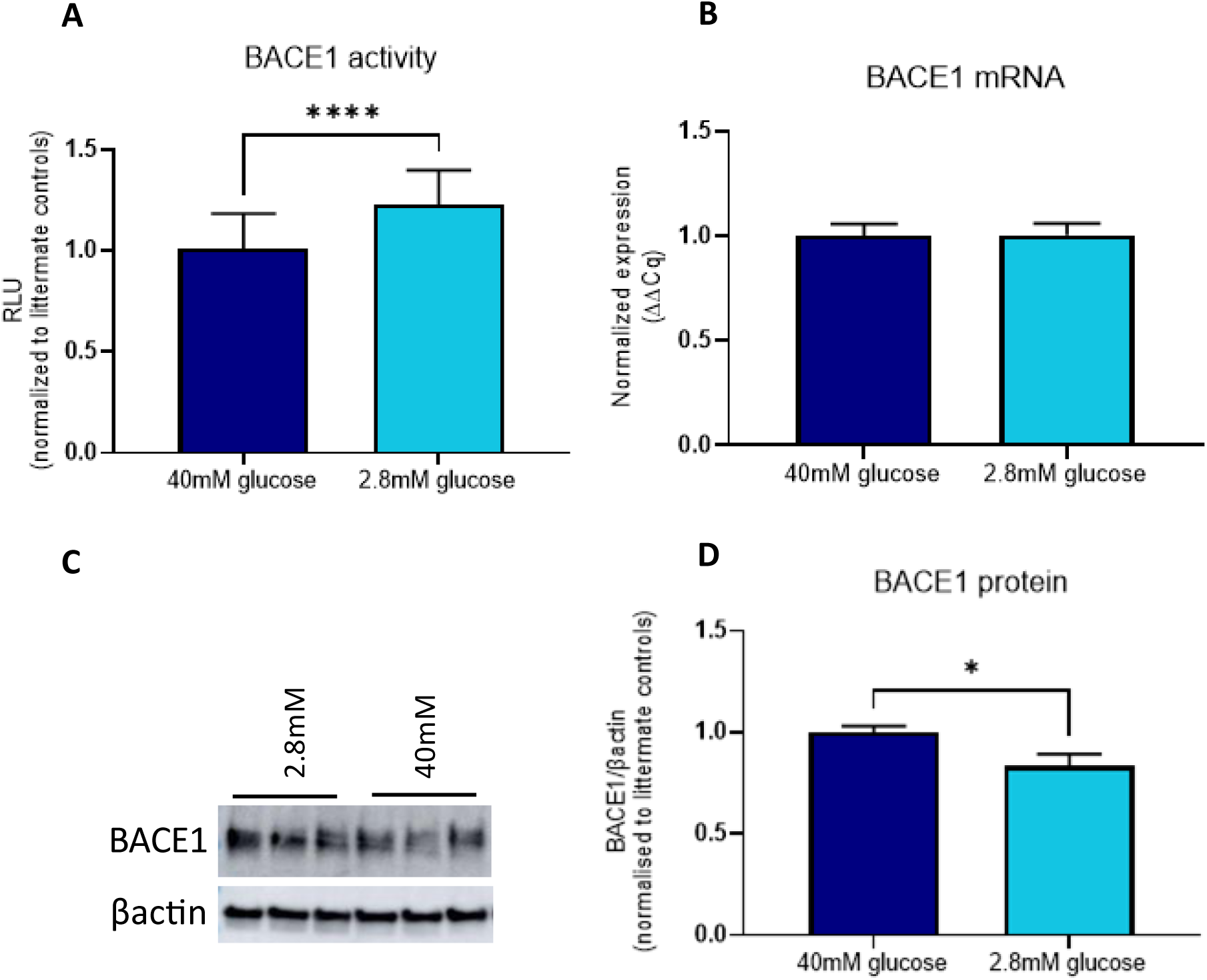
BACE1 activity is increased following hypoglycaemia. (a) WT OHSCs were treated with either 40mM glucose or 2.8mM for 7 days and lysates were harvested for BACE1 activity assay. There was a significant increase in activity (****p<0.0001) (n=17, paired t-test). (b) qPCR showed no significant change in BACE1 transcription (p=0.9692) (n=12; paired t-test). (c) yet there was a reduction in BACE1 protein following glucose treatment (*p=0.0154) (40mM n=14; 2.8mM n=13; unpaired t-test). Error bars = mean ± SEM.

### No indication of any change in γ-secretase activity

We also explored the possibility of alterations in γ-secretase activity due to altering glucose levels. Firstly, we sought to look for evidence of changes in PSEN1 and PSEN2, the genes responsible for encoding the catalytic components of γ-secretase, of which there are also dominant mutations in causing AD. However, qPCR analysis revealed no significant changes in the transcription of either of these genes (see Figure 5A-B). In further investigation, we examined downstream targets of Notch, which is another substrate cleaved by γ-secretase. qPCR analysis did not indicate any evidence of alterations in downstream targets for NOTCH signalling, including CCND1, Hes1, Hes5, Hey1, and Hey2 (Figure 5C-G).

**Figure 5.**
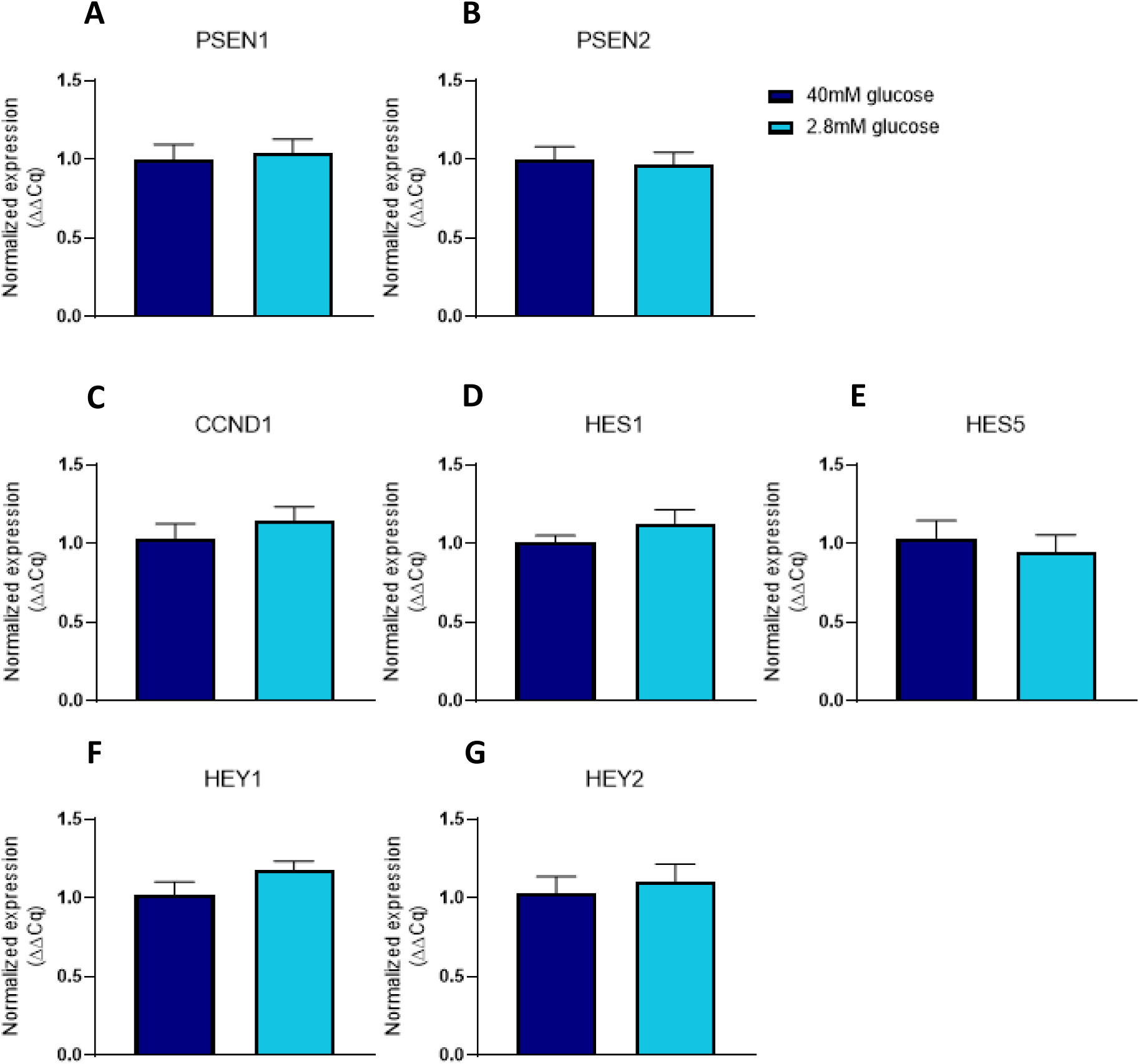
No detectable alterations in γ-secretase transcription or activity following glucose treatment. WT OHSCs were subject to 40mM or 2.8mM glucose concentrations for 7 days, and RNA collected for analysis. qPCR showed (a-b) no change in PSEN1 or PSEN2 levels (p= 0.7645 and p= 0.7882 respectively) (n=7, n=12 respectively; paired t-test), (c-g) nor was there any significant change in CCND1 (p= 0.2186), Hes1 (p=0.0929, Hes5 (p=0.6652), Hey1 (p=0.0636), or Hey 2 transcription (p=0.5140) (n=7; paired t-test). Error bars = mean ± SEM.

### No change in mRNA levels of enzymes involved in Aβ degradation

To pinpoint whether glucose-induced alterations in Aβ could also be driven by a decline in Aβ degradation, we investigated the effect of glucose concentration on several enzymes known to break down the peptide [56–61]. Following the same protocol, glucose levels were reduced to 2.8mM for 7 days, and mRNA was then extracted from WT slice tissue. qPCR analysis assessed mRNA levels of the following enzymes: angiotensin-converting enzyme (ACE), membrane metallo-endopeptidase (MME) (also known as neprilysin), insulin-degrading enzyme (IDE), cathepsin B 1 (CTSB1), cathepsin B 2 (CTSB2), and cathepsin D 1 (CTSD1). None of these genes exhibited a reduction in mRNA levels following glucose treatment (Figure 6A-F). It’s important to note that this list is not exhaustive, and the absence of a transcriptional change does not necessarily imply an absence of change in their activity.

**Figure 6.**
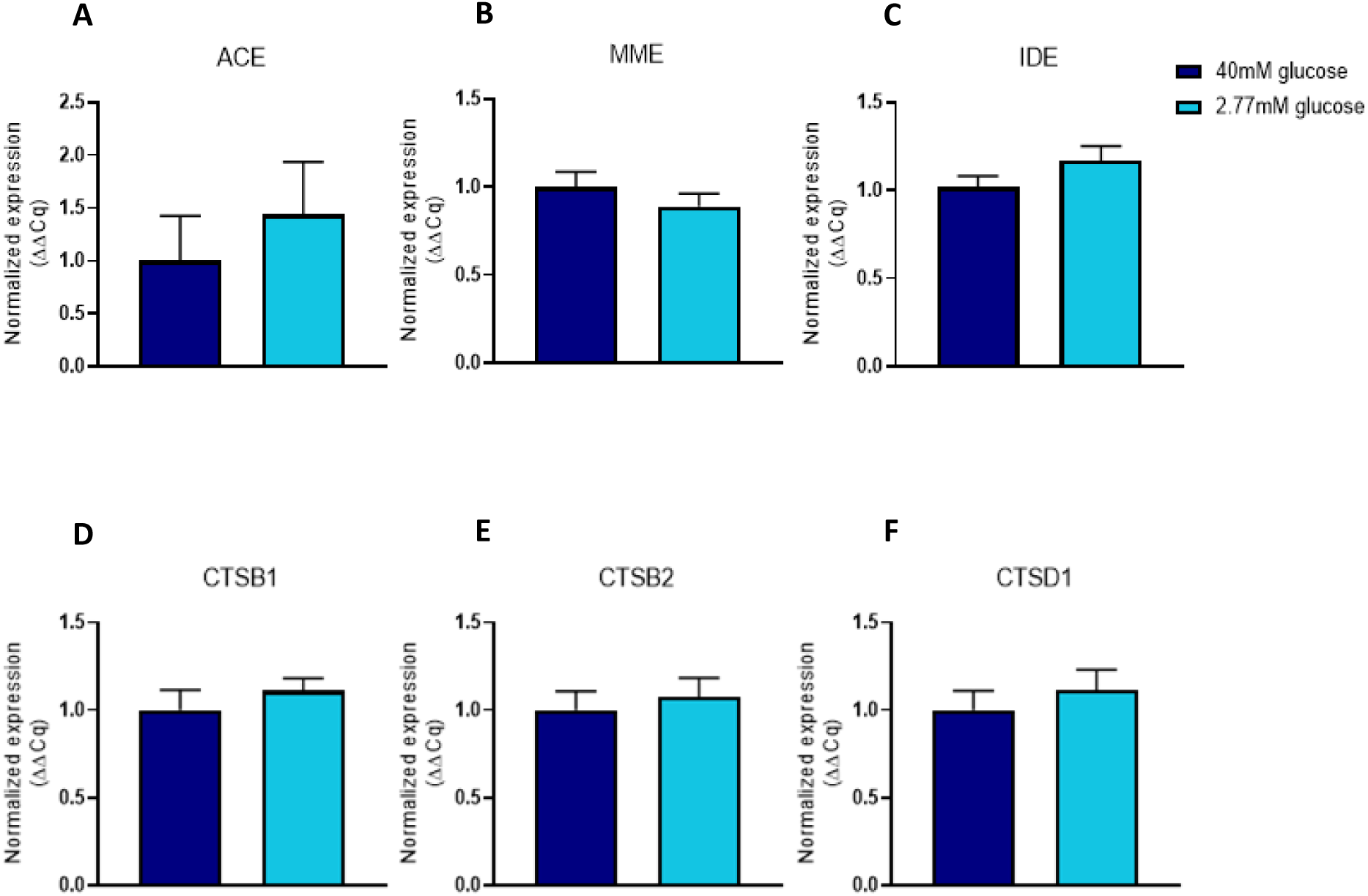
No transcriptional changes seen in key enzymes associated with Aβ degradation. WT OHSCs were cultured in either 40mM or 2.8mM glucose for 7 days and subsequently tissue harvested for qPCR. There was no significant change in any of the mRNA transcripts as follows: (a) ACE (p=0.5234) (n=12), (b) MME (p=0.1917) (n=12), (c) IDE (p=0.1360) (n=12), (d) CTSB1 (p=0.5193) (n=7), (e) CTSB2 (p=0.6172) (n=7), (f) CTSD1 (p=0.5255) (n=7) (paired t-test). Error bars = mean ± SEM.

### Pyruvate, lactate, and fructose 1,6 bisphosphate rescue low glucose induced change in Aβ

The brain’s reliance on glucose metabolism for energy entails that a substantial portion of glucose undergoes glycolysis, eventually entering the tricarboxylic acid (TCA) cycle and oxidative phosphorylation. To investigate the influence of the glycolysis primary end product on Aβ levels, OHSCs from WT mice underwent a 7 day treatment with 10mM pyruvate, in conjunction with varying glucose concentrations in the culture media. Figure 7A demonstrates that 10mM pyruvate effectively decreased Aβ levels in slices exposed 2.8mM glucose and had a trend for reduction with 40mM glucose. There was a statistical significance in the interaction between pyruvate and glucose.

**Figure 7.**
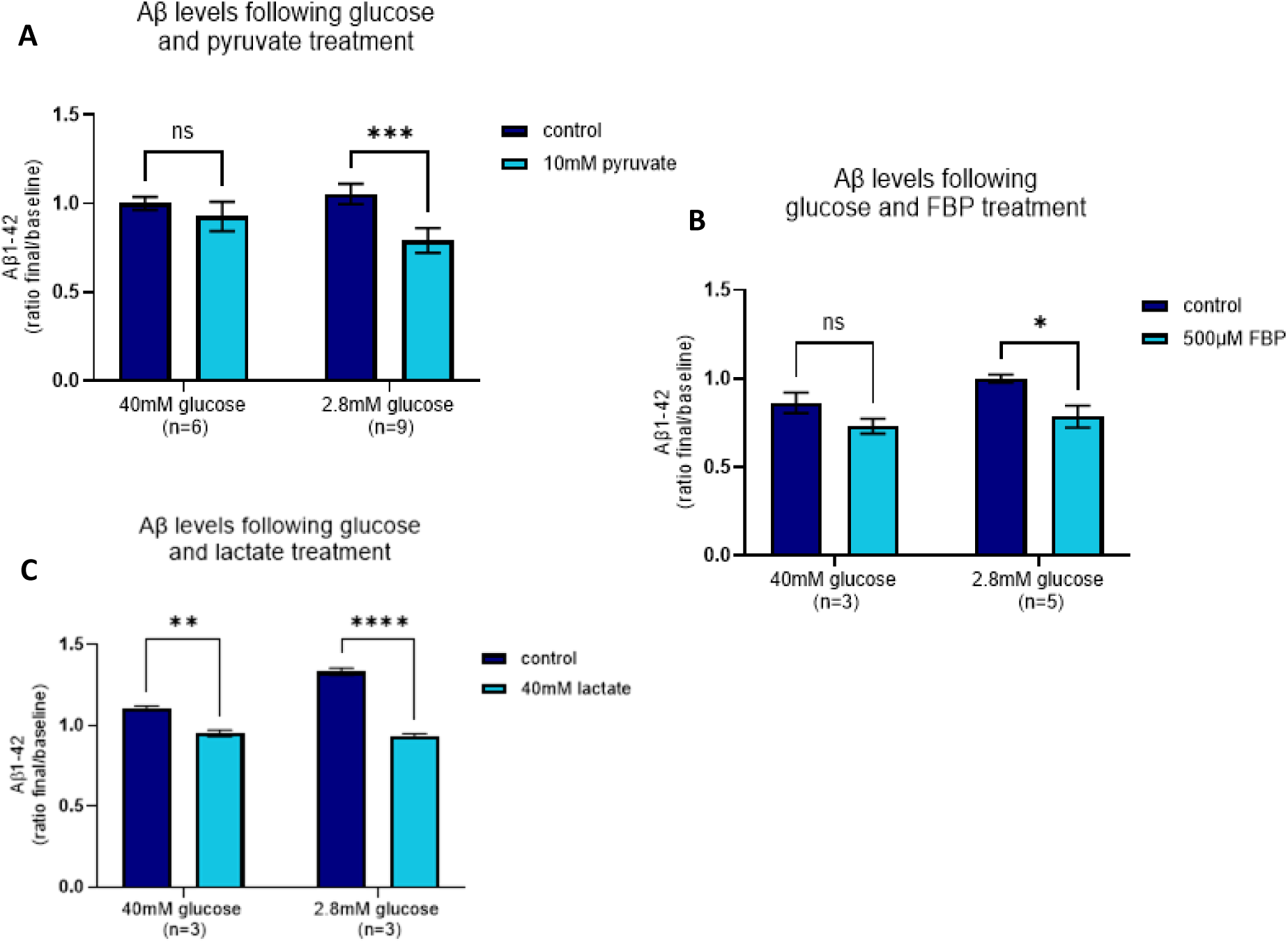
Treatment with pyruvate, FBP, and lactate reduce Aβ_1-42_ levels. WT OHSCs with either 40mM or 2.8mM glucose were co-treated with 10mM pyruvate for 7 days and medium collected for ELISA. (a) There was a significant reduction in Aβ_1-42_ when slices in low (2.8mM) glucose were treated with pyruvate (***p=0.0007) (n=9). A trend for a reduction was seen in 40mM glucose slices (p=0.4997) (n=6). Overall, there was a significant effect with pyruvate (**p=0.0018), a significant interaction between glucose: pyruvate (*p=0.0464), but no significance with glucose singularly (p=0.6324) (2way ANOVA with Sidak’s multiple comparisons test). (b) Similarly, slices were co-treated with 500uM FBP for 7 days. FBP treatment caused a significant decrease in Aβ_1-42_ levels when cultured in 2.8mM glucose (*p=0.0291) (n=5), and a trend was seen in 40mM glucose slices (p=0.2935) (n=3). Overall there was a significant effect due to FBP (*p=0.0155), but not glucose (p=0.1067), nor an interaction between the two (p=0.4501) (2way ANOVA with Sidak’s multiple comparisons test). (c) Finally, treating slices to 40mM lactate significantly reduced Aβ_1-42_ levels in both 40mM (**p=0.0017) and 2.8mM glucose (****p<0.0001). An overall significant effect was seen with lactate (****p<0.0001), glucose (***p=0.0004) and the interaction glucose: lactate (***p=0.001) (n=3; 2way ANOVA with Sidak’s multiple comparisons test). Error bars = mean ± SEM.

Fructose 1,6-bisphosphate (FBP) is positioned as the third metabolite in the glycolysis pathway and its production is considered the most crucial rate limiting step of glycolysis [62]. It has been proposed as a therapeutic intervention for conditions such as ischemia, epilepsy, and arthritis [63–66]. To examine whether FBP could counteract glucose-induced alterations in Aβ, WT OHSCs underwent a 7 day treatment with 500µM FBP, combined with varying glucose concentrations. This modest FBP concentration significantly reduced Aβ_1-42_ in the culture medium in combination with low glucose (2.8mM), and had a similar trend for a reduction of Aβ in 40mM glucose (Figure 7B).

Neurons in the brain and the OHSC system likely derive a portion of their energy through the lactate shuttle, and this reliance may undergo changes as part of a stress response, particularly during hypoglycaemic episodes [67]. The alteration in dependence on lactate could somehow lead to an increase in Aβ levels if lactate proves to be a less efficient energy source. Conversely, lactate might reduce Aβ levels by serving as an additional energy source. To investigate the impact of lactate on Aβ levels, slices exposed to both 40mM and 2.8mM glucose were co-treated with 40mM lactate for a 7-day period, and media samples were collected for ELISA. Similar to what is seen with pyruvate and FBP, lactate significantly reduced Aβ levels in both low glucose and control slices (Figure 7C). The reduction in the 2.8mM slices was slightly more pronounced, and there was a significant interaction between glucose and lactate, suggesting a potential rescuing effect.

## Discussion

There is a strong correlation between disrupted brain vasculature, glucose delivery and regulation, and the incidence of AD [13]. Individuals with vascular disease not only remain more susceptible to developing AD, but also the two pathologies share a significant number of risk factors. In this study, we demonstrate that lowering glucose levels in the culture media of OHSCs leads to an increase in Aβ production. This increase is associated with elevated BACE1 activity that has the potential to be causative. Importantly, this connection between glucose disruption and altered Aβ occurs in WT tissue, without resorting to genetic manipulation and thus represents one possible mechanism in sporadic AD that elevates Aβ.

Our findings reveal that reducing glucose concentrations from a normal 40mM to 2.8mM in the culture medium results in elevated levels of Aβ_1-42_ and Aβ_1-40_ excreted by WT OHSCs. This increase is not confined to the medium, as we observe a parallel rise in Aβ_1-42_ within the tissue. While energy inhibition through the form of insulin or glycolysis inhibitors have shown an increase in Aβ in APP transgenic animals [68], this is the first time it has been shown to induce a change in Aβ in a WT model. The co-occurrence of elevated Aβ_1-42_ and Aβ_1-40_ suggests that the shift is not due to alterations in the ratio of these two Aβ species, but we cannot exclude changes in other isoforms, as the length of Aβ varies between 37-49 amino acids. A similar increase in Aβ levels in human SH-SY5Y cells and OHSCs derived from the TgCRND8 model of human APP indicates a non-species-specific response and a neuronal origin.

Moreover, we investigated hypoxia, which is relevant in scenarios involving diminished brain vasculature, and found no substantial impact of this upon Aβ levels, either in isolation or when combined with hypoglycaemia. This stands in contrast to earlier studies indicating that hypoxia can trigger an increase in Aβ levels in APP transgenic mice and SH-SY5Y cells overexpressing Swedish APP, albeit those studies focused on shorter yet more severe bursts of hypoxia [69,70]. It has been demonstrated that known AD mutations can render astrocytes more susceptible to ischemic events [71], potentially explaining the limited effect observed in WT slices.

To determine what was upstream of the glucose-induced effect upon Aβ, we first examined APP levels. O’Connor and colleagues have previously reported a significant *decrease* in APP mRNA (despite increasing Aβ_40_ production) with insulin treatment or glycolysis inhibition. Yet, APP protein is reportedly *increased* in Tg2576 animals following energy inhibition [68]. Our investigation into APP levels following glucose reduction showed unaltered APP transcription and translation in WT and TgCRND8 OHSCs. This does not rule out any changes in post-translational regulation such as glycosylation, phosphorylation or sulfation [72]. An alternative rationale for the rise in Aβ levels due to hypoglycaemia could involve a potential shift in the location of APP. APP processing is dependent in part upon its trafficking and sub-cellular location, and enzymatic cleavage occurs within the extracellular membrane and endosomes [73–75]. To explore this possibility, co-staining with APP and markers for endosomes could be employed to examine whether there are any alterations in the protein’s cellular localization.

Next, we investigated BACE1, the first enzyme in the amyloidogenic pathway responsible for the cleavage of APP into Aβ. We show that BACE1 activity exhibited a significant increase without corresponding alterations in BACE1 mRNA or protein levels. This partially contrasts with studies demonstrating increased BACE1 protein levels in glucose deprivation in WT animals and HEK283 cells [76], and in Tg2576 mice subjected to insulin treatment and glycolysis inhibition [68]. Yet, BACE1 undergoes many post-translational modifications which have the ability to affect its activity without an increase in its expression [39]. Indeed, the slight lowering of BACE1 protein that we observed could represent a compensatory response for increased activity caused by a post-translational step. Similar to APP, it would be interesting to look further into the cellular localization of BACE1, as it can heavily influence its activity and Aβ production. Notably, our results suggests a potential avenue of exploration of a direct link between vascular disfunction and β-secretase activation in WT tissue.

Presenilins make up the catalytic component of γ-secretase, which further cleaves APP into Aβ [77]. Similar to APP mutations and ischemia, presenilin mutants have displayed an elevated susceptibility to glucose deprivation [78]. Subsequent analysis of hypoglycaemia treatment in WT slices explored γ-secretase and its targets, revealing no impact of glucose levels on PSEN1 or PSEN2 transcription. In an effort to determine whether there was any noticeable effect upon γ-secretase activity, we looked at markers of NOTCH signalling given that this protein serves as both a target of, and its pathway activated by, γ-secretase [79–81]. qPCR following glucose treatment showed that downstream targets of NOTCH were not conclusively affected. Similarly, we looked for transcriptional changes in enzymes involved in Aβ degradation, including ACE [58], IDE [23,56], the cathepsins [60,61,82,83] and MME, also known as neprilysin [57]. Once again, there was no indication of alterations in mRNA for these enzymes, although, this does not rule out a change in their activity.

Finally, several glycolysis metabolites and potential fuel substitutes were tested for their ability to mitigate the glucose-induced increase in Aβ. Pyruvate, FBP, and lactate all demonstrated the capacity to reduce Aβ levels in WT OHSCs. A pyruvate and lactate-induced decrease in Aβ occurred under both normal and hypoglycaemic glucose conditions and showed a significant interaction between the compounds and glucose treatment. In contrast, FBP only significantly decreased Aβ in 2.8m glucose medium. The means by which these compounds decreased Aβ is unclear will be an important topic for further investigation. FBP is of particular interest as it its currently being suggested as a potential therapy for epilepsy and ischemia [63,64,66]. While glucose failed to have a significant effect upon Aβ production in the pyruvate and FBP experiments, this is likely due to the fact that 40mM and 2.8mM glucose OHSCs were not cultured from the same animals. This made them subject to intra-pup variability in Aβ levels, requiring a higher n-number to reach significance. Overall, the ability of these substrates to directly influence Aβ in WT OHSCs underscores the value of these cultures in dementia research.

## Conclusions

In summary, our findings demonstrate the ability to induce an increase in Aβ production in WT and APP transgenic brain tissue through the manipulation of glucose availability in an *ex vivo* system. This coincides with a concurrent rise in BACE1 activity. Furthermore, the introduction of additional metabolites exhibited the potential to decrease Aβ levels. Given that a majority of AD cases are sporadic, it becomes crucial to comprehend the development of amyloid pathology in the absence of causal mutations in APP or the presenilins. The ability to first elevate Aβ in WT tissue, and subsequently alleviate such effects as we report here, provides valuable insights into how future therapeutics aimed at preventing or slowing down the progression of the disease could be developed.

## List of abbreviations

Aβ: amyloid-β
ACE: angiotensin-converting enzyme
AD: Alzheimer’s disease
APP: amyloid precursor protein
BACE1: β-site APP Cleaving Enzyme 1
CTSB1: cathepsin B 1
CTSB2: cathepsin B 2
CTSD1: cathepsin D 1
FBP: fructose 1,6-bisphosphate
IDE: insulin-degrading enzyme
MME: membrane metallo-endopeptidase
NFTs: neurofibrillary tangles
OHSC: organotypic hippocampal slice culture
WT: wildtype

## Declarations

### Ethical approval

Animal work was approved by Cambridge and performed in accordance with the Animals (Scientific Procedures) Act 1986 under Project licenses 70/7620 and P98A03BF9.

### Consent for publication

Not applicable

### Availability of data and material

The datasets used and/or analysed during the current study available from the corresponding author on reasonable request.

### Author’s Contributions

Study concept and design: OS, CD and MC. Data acquisition: OS and RH. Statistical analysis: OS. Analysis and interpretation of the data: OS and MC. OS, RH, CD and MC co-wrote the manuscript. All authors read and approved the final manuscript.

## Acknowledgements

Not applicable

## Author’s information

Not applicable

---

**Supplemental table 1.**
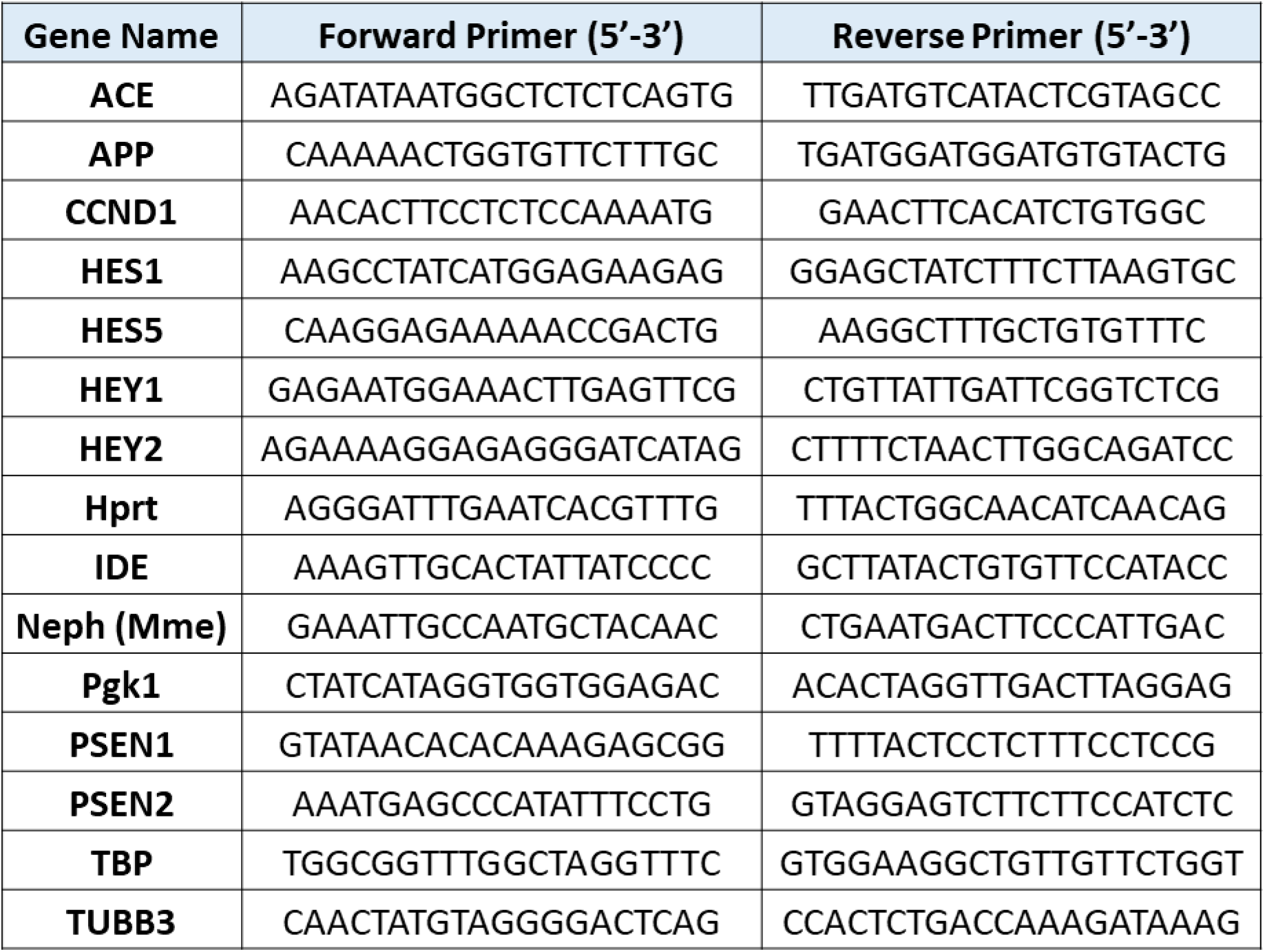
List of primers for qPCR analysis.

**Supplemental Figure 1.**
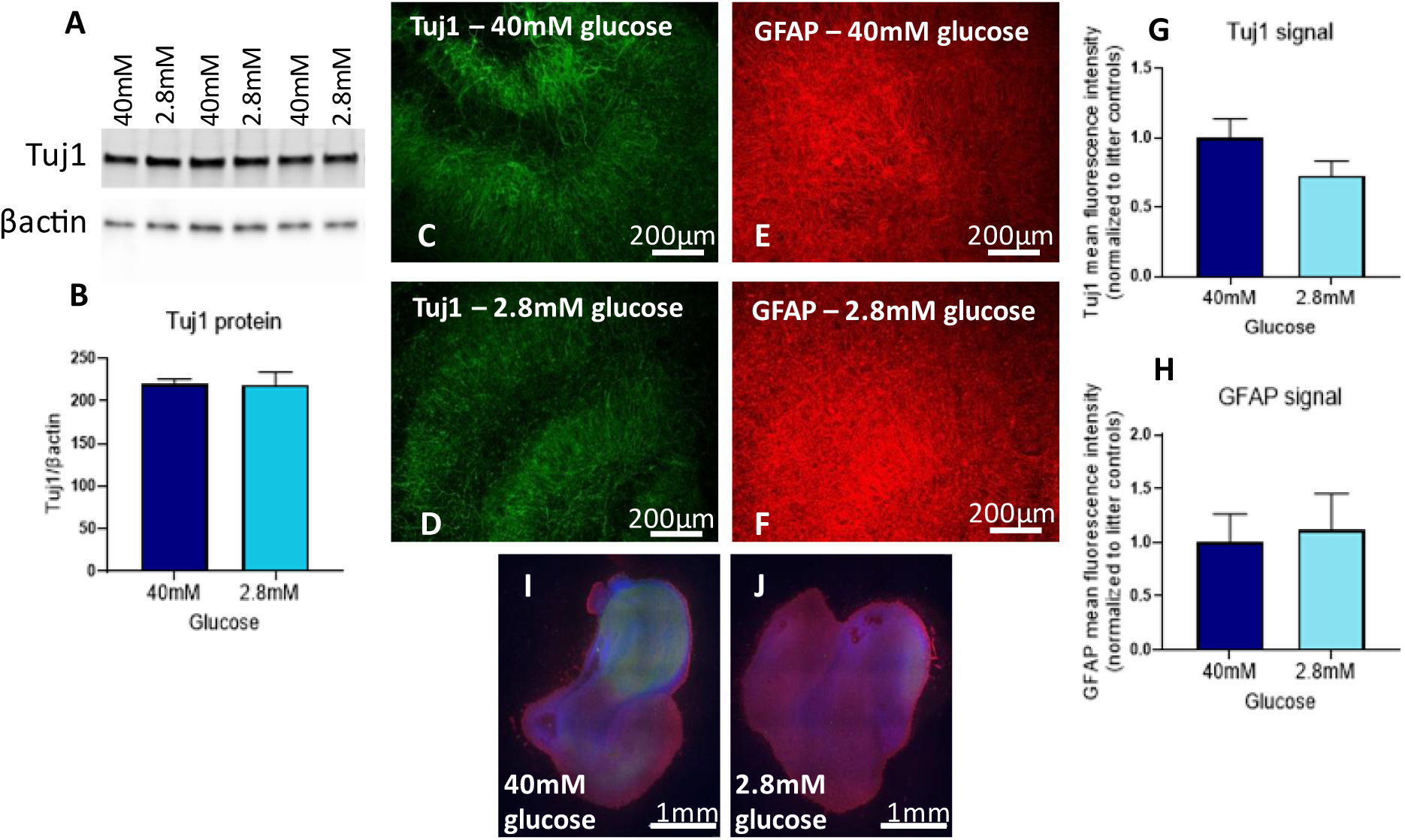
Reducing glucose in the culture medium does not affect neuronal viability after 7 days, but is toxic by 21 days. WT OSCSs were cultured in either 40mM or 2.8mM glucose for 7 days. (a) Slices were harvested for immunoblotting and showed no significant change in TUJ1 protein (p=0.9448) (n=4; paired t-test). Similarly, (c,d,g) slices stained for TUJ1 showed no significant change (0.2242) (n=4), and (e,f,h) stained for GFAP showed no significant change (p 0.6468) (n=4) (paired t-test). (i-j) By 21 days of glucose treatment, immunostaining with TUJ1 showed a gross reduction in signal. Error bars = mean ± SEM.

